# Time-resolved cryo-EM reveals conformational trajectory of allosteric activation in isocitrate lyase

**DOI:** 10.64898/2026.04.08.716820

**Authors:** Jamie Taka, James Jung, Shuning Guo, Wanting Jiao, Brooke X. C. Kwai, Luiz Pedro S de Carvalho, Matthew McNeil, Evelyn Yu-Wen Huang, Zhiheng Yu, Ivanhoe K. H. Leung, Ghader Bashiri

**Affiliations:** School of Biological Sciences, The University of Auckland, Auckland, Private Bag 92019, New Zealand; Janelia Research Campus, Howard Hughes Medical Institute, Ashburn, VA 20147; Ferrier Research Institute, Victoria University of Wellington, Wellington 6140, New Zealand; School of Chemical Sciences, The University of Auckland, Auckland, Private Bag 92019, New Zealand; Department of Chemistry, The Herbert Wertheim UF Scripp Institute of Biomedical Innovation and Technology, Jupiter, FL, USA; Department of Biochemistry, University of Otago, Dunedin, New Zealand; Bio21 Molecular Science and Biotechnology Institute, School of Chemistry, University of Melbourne, Parkville, Victoria 3052, Australia

## Abstract

Isocitrate lyase 2 (ICL2) from *Mycobacterium tuberculosis* undergoes dramatic conformational rearrangements upon binding to the allosteric effector acetyl-CoA. Time-resolved cryo-EM captured conformational states along the ICL2 activation trajectory, revealing how acetyl-CoA binding at the allosteric sites leads to asymmetric, half-of-site activity at the catalytic centres. These findings support a conformational selection model of allostery, whereby acetyl-CoA binding shifts the pre-existing equilibrium towards an active state of the enzyme.

## Main

Isocitrate lyase 2 (ICL2) acts as a gatekeeper enzyme in *Mycobacterium tuberculosis* (*Mtb*), regulating central carbon metabolism under various physiological and pathological conditions^1–4^. This regulation occurs through allosteric activation of the isocitrate lyase and methylisocitrate lyase activities of *Mtb*-ICL2 upon binding of acetyl-CoA or propionyl-CoA^1^, which are end-products of multiple catabolic pathways, including glycolysis, pyruvate/lactate, and lipid metabolism. ICL2, together with ICL1, is essential for *Mtb* survival during the persistent stage of infection^5–7^.

*Mtb*-ICL2 forms a tetramer, in which the N-terminal domains create a central catalytic core, while the C-terminal domains form dimers at either end in the apo state (inactive dimers) and interact with the helical subdomain (Figure 1.a). Acetyl-CoA binding to the C-terminal domains results in a dramatic rearrangement, wherein the C-terminal domains shift toward the catalytic core, forming new dimer pairs (active dimers). This acetyl-CoA stabilised conformation increases catalytic efficiency by 50-fold^1^. However, the mechanism linking acetyl-CoA binding to these structural and functional changes remains unclear. Here, we combine time-resolved cryo-electron microscopy (cryo-EM), small-angle X-ray scattering (SAXS), isothermal titration calorimetry (ITC), and kinetic analysis to elucidate this mechanism.

**Figure 1.**
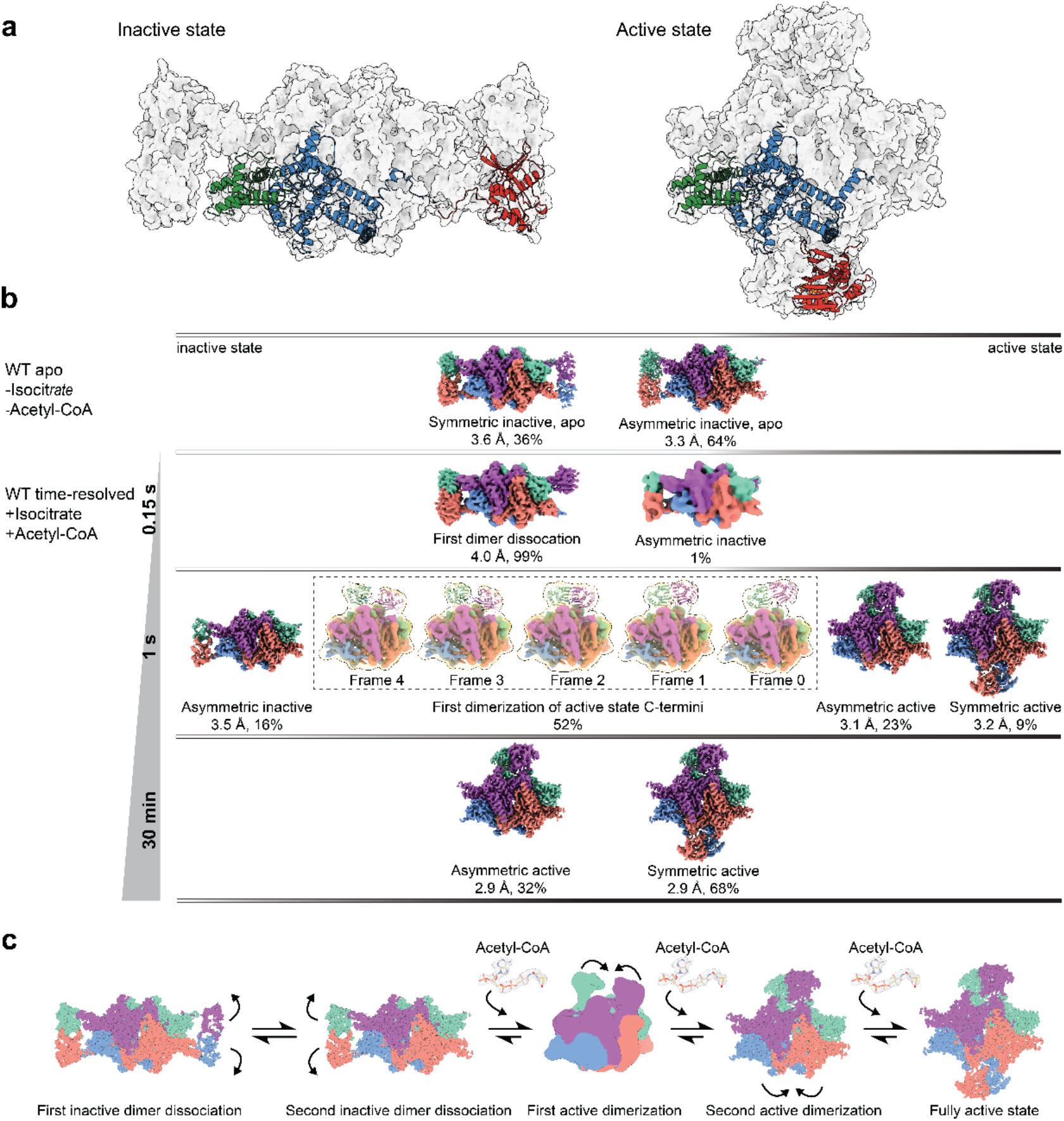
Time-resolved cryo-EM analysis of *Mtb-*ICL2 activation by acetyl-CoA. **a**, Crystal structure of *Mtb-*ICL2 in the absence of acetyl-CoA (left, PDB code: 6EDW) and bound by acetyl-CoA (right, PDB code: 6EE1). The N-terminal domain (blue) contains the active site, while the C-terminal domain (red) binds acetyl-CoA and is connected to the N-terminal domain via a flexible linker. The helical subdomain (green) stabilises the inactive conformation by interacting with the linker of the neighbouring monomer. **b**, Cryo-EM maps of *Mtb-*ICL2 in the apo state, or after addition of acetyl-CoA and substrates (isocitrate/Mg^+2^) at 150 msec, 1 sec, and 30 mins. Percentages indicate the fraction of particles contributing to each 3D reconstruction. **c**, Proposed model for *Mtb-*ICL2 activation upon acetyl-CoA binding.

We applied time-resolved cryo-EM to capture conformational states of *Mtb*-ICL2 during activation in the presence of the substrate (isocitrate/Mg^2+^) and the allosteric effector acetyl-CoA (Figure 1.b, Figure S1-4). We examined *Mtb*-ICL2 at three time points: 0.15 sec, 1 sec, and 30 min. The earliest time point (0.15 sec) was achieved using a microfluidics mixer-sprayer-plunger apparatus, which allowed rapid mixing of enzyme, substrate, and acetyl-CoA immediately before vitrification. For the 1 sec and 30 min time points, samples were manually mixed on grids and flash-frozen in liquid ethane.

At 0.15 sec, *Mtb*-ICL2 predominantly (99%) adopts an inactive state in which one pair of C-terminal dimers has dissociated (Figure 1.b, Figure S2) with one of the C-terminal domains lacking any discernible density. This indicates that the dissociated C-terminal dimer likely represents the conformation that favours acetyl-CoA binding. Notably, this dissociation is essential for acetyl-CoA binding, since an engineered inactive variant (D377C/L601C, F381C/T604C) in which two disulfide bridges lock the helical subdomain and linker, neither binds acetyl-CoA (Figure S5) nor exhibits catalytic activity (Figure S6). Further, the isolated C-terminal domain is monomeric in solution^8^ and binds acetyl-CoA with a dissociation constant of 50 μM (Figure S5). By 1 sec, while both inactive (16%) and active (32%) conformations are present, over half the population (52%) represents intermediate stages of active dimer formation, with monomers forming transient contacts with the N-terminal core before stabilising as active dimers (Figure 1.b, Figure S3). At 30 min, only active conformations remain, with both symmetric (68%) and asymmetric (32%) states present (Figure 1.b, Figure S4). The symmetric active state closely resembles the acetyl-CoA-bound crystal structure (PDB code: 6EE1) with an RMSD of 1.7 Å for all equivalent atoms, whereas the asymmetric active state was not observed in crystal structures and shows density for only one active C-terminal dimer. Interestingly, acetyl-CoA binding shows positive cooperativity (Hill number of 1.6, Figure S7), with the initial binding enhancing the affinity for subsequent binding events in the enzyme. Together, this supports a conformational selection model of allostery, whereby acetyl-CoA binding shifts the pre-existing equilibrium towards an active state of the enzyme. These data provide unprecedented insight into the structural trajectory of ICL2 activation stabilised by acetyl-CoA binding (Figure 1.c).

Our time-resolved cryo-EM further reveals a dynamic feature of the enzyme that provides a plausible mechanism linking acetyl-CoA binding to conformational changes and catalytic activation. 3D variability analysis of active *Mtb*-ICL2 states reveals a 5° pivot of each active C-terminal dimer around a fulcrum formed by the Q635-E640 loop of both monomers, producing a 4 Å displacement at the furthest points of the dimer (Figure 2.a, Video S1-2). The flexible linker is more ordered when the C-terminal domain is closest to the N-terminal core and disordered in the opposing monomer (Figure 2.b). In this conformation, the ordered linker lies adjacent to the active site loop, likely stabilising the ‘closed’ catalytic conformation, while the opposing loop remains ‘open’ for substrate entry or product release. This ‘seesaw’ mechanism results in alternating half-of-site reactivity, with only one active site per dimer catalytically engaged at a time, a regulatory feature that has been observed in several multimeric enzymes^9–11^.

**Figure 2.**
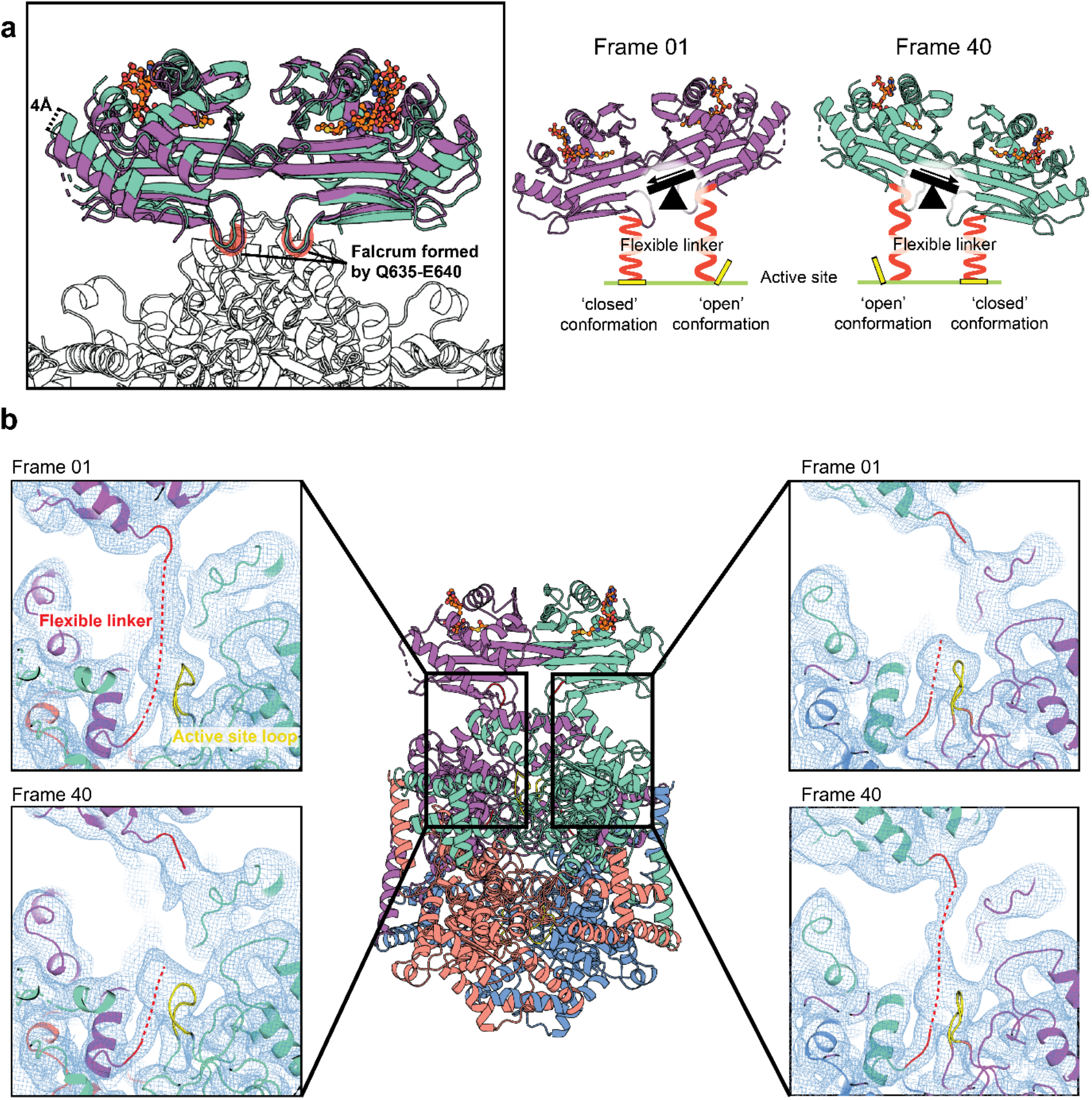
Cryo-EM analysis reveals a seesaw mechanism regulating *Mtb-*ICL2 activity. **a**, 40 frames from 3D variability analysis of *Mtb-*ICL2 in the active conformation illustrate a rocking motion of the C-terminal dimer. Comparison of these frames shows a 5° pivot of the C-terminal dimer between the first and last frames. In the proposed ‘seesaw’ mechanism, the rocking motion of the active C-terminal dimer results in the conformational change of the linker connecting each C-terminal monomer to the N-terminal core. In one monomer, the linker is positioned adjacent to the active site loop, stabilising a ‘closed’ active site loop conformation that promotes catalysis. In the opposite monomer, the linker is less ordered, resulting in an ‘open’ loop conformation that permits substrate entry or product release. **b**, Cryo-EM density for the active site (yellow) and flexible linker (red) in frames 1 and 40 of the active conformation.

To test this model, we generated two sets of *Mtb*-ICL2 variants. First, our model places the flexible linker adjacent to the active site loop (^213^KKCGH^217^). We mutated residues within the flexible linker to generate a Q597E/H598E variant that enhances isocitrate lyase activity to a level comparable to the activated wild-type enzyme in the absence of acetyl-CoA (Figure S8). This is presumably due to enhanced charge-charge interactions between the linker and the active site loop. Second, we created a stabilised active variant of *Mtb*-ICL2 (E640C/N731C/E733C) in which three disulfide bonds lock the C-terminal domains in an active dimer conformation^8^. The crystal structure of the isolated C-terminal domain confirms the position of the engineered disulfide bonds (Figure S9). These disulfide bonds did not alter acetyl-CoA binding in either the full-length enzyme (1.9 *vs* 2.7 μM, wild-type *vs* active variant, respectively) or the isolated C-terminal dimer (1.7 μM) (Figure S10). Furthermore, SAXS profiles show that the full-length active variant adopts an active conformation even in the absence of acetyl-CoA (Figure S11). Supporting our hypothesis, kinetic analysis revealed that the turnover number (*k*_cat_) for the active variant was double that of the activated wild-type (1.2 s^-1^ *vs* 2.5 s^-1^, wildtype *vs* active variant, respectively, Table S1). This suggests that all four active sites are catalytically proficient, compared to two in the wild-type, which comes at a cost of reduced *K*_M_ for the substrate isocitrate (1.2 mM *vs* 0.7 mM, wildtype *vs* active variant, respectively, Table S1).

In summary, time-resolved cryo-EM enabled the direct visualisation of conformational states during the allosteric activation of *Mtb*-ICL2, linking effector binding to the catalytic mechanism. *Mtb*-ICL2 demonstrates the applicability of time-resolved cryo-EM to metabolite-driven enzyme allostery, extending beyond the highly modular allosteric systems recently reported^12–16^. In *Mtb*-ICL2, acetyl-CoA binding shifts a weakly populated conformational equilibrium toward the active state, providing structural support for a conformational selection model of allostery^17^. Together, these insights advance our understanding of protein allostery that underpins enzyme function and regulation.

