## Supplementary Information for "Time-resolved cryo-EM reveals conformational trajectory of allosteric activation in isocitrate lyase"

### Materials and Methods

#### Expression and purification

The open reading frame encoding the wild-type ICL2 of *Mycobacterium tuberculosis* CDC 1551 and all ICL2 variants were cloned into pYUB28b[1,2]. Recombinant ICL2 was expressed in *Escherichia coli* BL21 (DE3) LOBSTR cells[3] transformed with the pGro7 plasmid (Takara Bio Inc,) expressing GroEL/GroES chaperones under the *araB* promoter. ICL2 constructs were expressed in Terrific Broth media containing 50 µg/mL of hygromycin B and 34 µg/mL of chloramphenicol at 18°C. Expression was induced by the addition of 1 mM isopropyl β-D-1-thiogalactopyranoside (IPTG) and L-arabinose (final concentration 0.5 g/L) to log phase cultures prior to subsequent growth overnight at 18°C. Cells were lysed by cell disruption in 20 mM Tris-HCl (pH 8.0), 300 mM NaCl, 10% (v/v) glycerol, and 0.1% (v/v) Triton X-100. The hexahistidine-tagged ICL2 was purified by immobilised metal affinity chromatography and size exclusion chromatography using a HiLoad 16/600 Superose 6 pg column (Cytiva) equilibrated in buffer containing 10 mM Tris-HCl (pH 8.0) and 150 mM NaCl.

#### Enzyme kinetics

Enzyme kinetics were measured using a phenylhydrazine-coupled UV/Vis assay[2] in a 96-well UV-Star® microplate (Greiner bio-one) with a SpectraMax iD3 plate reader set to 20°C. All experiments were undertaken in buffer containing 50 mM Tris (pH 8.0), 5 mM $MgCl_2$, 150 mM NaCl, 10 mM phenylhydrazine, and 200 nM enzyme. For samples containing acetyl-CoA, a final concentration of 25 µM acetyl-CoA (Roche) was added. Reactions were initiated by the addition of DL-isocitrate (Thermo Fisher Scientific). The product of isocitrate lyase, glyoxylate, reacts with phenylhydrazine to form a phenylhydrazone adduct that is measured spectrophotometrically at 324 nm. The absorbance was converted to concentration using a calibration curve constructed using known concentrations of glyoxylate and phenylhydrazine in the same reaction buffer. Initial rates were obtained from the regions where the substrate turnover was <10%. Nonlinear regression was used to fit the data to determine the Michaelis-Menten kinetics (Equation 1). A summary of the enzyme kinetics is found in Table S1.

Acetyl-CoA binding cooperativity was determined using the same phenylhydrazine-coupled UV/Vis assay with 200 nM ICL2, 1 mM DL-isocitrate, and a two-fold concentration series of acetyl-CoA (0 - 80 µM). Initial rates were obtained from the regions where the substrate turnover was <10%. Nonlinear regression was used to fit the data to the Hill equation to determine the Hill coefficient (Equation 2).

Equation 1:

$$v = \frac{Vmax[S]}{Km + [S]}$$

[S]: Substrate concentration

Vmax: Maximum velocity

Km: Michaelis Constant

$v$: Initial velocity

Equation 2:

$$v = \min + \frac{max - min}{1 + \left(\frac{EC_{50}}{[x]}\right) Hill}$$

[x]: Acetyl-CoA concentration

min: Minimum response (i.e., no acetyl-CoA)

max: Maximum response

$EC_{50}$: Concentration of ligand to produce half the maximum response

Hill: Hill coefficient

**Small-angle X-ray scattering**

ICL2 constructs were dialysed against buffer containing 20 mM Tris pH 8.0, 150 mM NaCl and 5% glycerol (v/v). Proteins were made to 2.5–5 mg/mL and, where appropriate, incubated with 0.5 mM acetyl-CoA before SAXS analysis. SAXS data were collected on the Australian Synchrotron BioSAXS beamline. The samples were mounted in the CoFlow autosampler at ~293 K for capillary flow data acquisition[4]. Data was processed using in-house data-reduction software developed by the Australian Synchrotron and the ATSAS software package[5].

**Isothermal titration calorimetry**

ICL2 constructs were dialysed against phosphate buffer containing 10 mM sodium phosphate pH 8.0, and 150 mM NaCl. Acetyl-CoA was dissolved directly in phosphate buffer. Acetyl-CoA was titrated into a sample cell containing ICL2 maintained at 20°C. Each injection was spaced 300 s apart to allow a baseline to be reached, and the reference power was set to 10 µCal $sec^{-1}$. The heats of dilution of acetyl-CoA were measured by titrating acetyl-CoA into phosphate buffer. The calorimetric data were analysed using the Origin® software package. The heats of dilution of acetyl-CoA were removed from the binding data. The full-length wild-type ICL2 construct was used at sample cell concentrations of 60 µM with 1000 µM acetyl-CoA. The full-length active variant (E640C/N731C/E733C) and inactive variant (D377C/L601C, F381C/T604C) were used at sample cell concentrations of 50 µM with 500 µM acetyl-CoA. The ICL2 isolated C-terminal domain (residues 601-766) with engineered disulfide bonds (E640C/N731C/E733C) was used at a sample cell concentration of 30 µM with 250 µM acetyl-CoA. The wild-type isolated C-terminal domain was used at a sample cell concentration of 550 µM with 7000 µM acetyl-CoA.

### Protein X-ray crystallography

Crystals of the isolated C-terminal domain with engineered disulfide bonds (E640C/N731C/E733C) were obtained by sitting drop vapour diffusion with 8 mg/mL protein and 1 mM acetyl-CoA. Crystals formed in drops containing a 1:1 ratio of protein and a crystallisation condition with 0.1 M Bis-Tris pH 5.5, 25% w/v PEG 3350. Diffraction data were collected using the MX1 beamlines at the Australian Synchrotron. All datasets were indexed and processed using XDS[6], and scaled with AIMLESS[7] from the CCP4 programme suite[8]. Complete data collection and processing statistics are given in Table S2. The structure was solved by molecular replacement with Phaser[9] using the C-terminal dimer (chain B&D residue 601-766) of the full-length *Mtb*-ICL2 crystal structure (PDB ID: 6EE1)[2]. The initial model was further refined using REFMAC5[10] and manually built with COOT[11].

### Cryo-EM sample preparation

The apo state wild-type ICL2 samples were concentrated to 3.35 mg/mL. The cryo-EM grids were prepared using Chameleon (SPT Labtech) on self-wicking grids in two-stripe mode at ~54 ms wicking time[12]. The self-wicking grids were glow-discharged for 30 s at 12 mA internally prior to sample dispensing. The 1 s and 30 min time-resolved wild-type ICL2 samples were concentrated to 1.5 mg/mL. The samples were preincubated with 0.5 mM DL-isocitrate for 30 min at 21°C. The grids were first applied with 0.4 µL 5 mM acetyl-CoA, then 3.6 µL wild-type ICL2, immediately followed by blotting single-sided using Leica EM GP2 at 15°C and 100% relative humidity level for 1 s, resulting in 6 s overall mixing time before trapping by vitrification in liquid ethane. Quantifoil R2/1, 400 mesh holey carbon grids (Quantifoil Micro Tools) were glow-discharged for 30 s with a 15 mA current under an evacuated air pressure of 0.39 mbar in a PELCO easiGlow system (Ted Pella Inc). The 0.15 s time-resolved dataset wild-type ICL2 sample was concentrated to 1.7 mg/mL. A polydimethylsiloxane (PDMS) microfluidic chip-based sample mixer-sprayer time-resolved plunging instrument similar to the device described by Joachim Frank and Stephen P. Muench was used, but uniquely in combination with self-wicking grids glow-discharged as described above prior to sample dispensing[13,14]. 0.5 mM acetyl-CoA and 0.5 mM DL-isocitrate were sprayed coincidentally with wild-type ICL2 on the self-wicking grids for an average mixing time of 0.15 s before trapping by vitrification in liquid ethane.

### Cryo-EM data collection

Movies of ICL2 particles embedded in vitreous ice were collected at liquid nitrogen temperature using Titan Krios transmission electron microscopes (Thermo Fisher Scientific). The 1 s time-resolved and apo state wild-type ICL2 datasets were collected on Janelia Krios3 equipped with an Extreme Brightness Cold Field Emission Gun (E-CFEG, Thermo Fisher Scientific), a Selectris X energy filter (Thermo Fisher Scientific) with a 5 eV-wide energy slit and a Falcon4i direct electron detector (Thermo Fisher Scientific). The end point wild-type ICL2 dataset was collected on Janelia Krios2 microscope equipped with a high-brightness Schottky FEG (X-FEG, Thermo Fisher Scientific) operated at

300kV, a Gatan Imaging Filter (GIF) with a 10 eV-wide energy slit and a K3 direct electron detector (Gatan). The 0.15 s time-resolved dataset was collected on Janelia Krios1 microscope equipped with a CETCOR spherical aberration (*C*s) corrector (Corrected Electron Optical Systems GmbH) with a *C*s level of 0.01 mm, a Gatan BioContinuum HD energy filter operated with a 10 eV-wide energy slit and a K3 direct electron detector (Gatan Inc). The K3 movies were recorded in correlative double sampling (CDS) mode with a binning of 0.5, and later binned two-fold. The Falcon4i movies were recorded with binning 1 in Electron Event Representation (EER) format. Data was collected at a nominal magnification of 81,000–165,000 X with calibrated pixel sizes between 0.743–0.857 Å. A defocus range of -0.8 – -2.6 µm was applied. The total electron dose of each movie was 60 $e^-/Å^2$. Detector gain reference images were collected at the start of each data collection session, but were not applied during data collection. All data was collected using a semi-automated workflow and customised scripts in SerialEM. The data collection parameters are in Table S3.

### Cryo-EM data processing

Beam-induced motions of particles were corrected using Unblur of *cis*TEM2[15,16]. Contrast transfer function (CTF) parameters were estimated from sums of three movie frames using *cis*TEM2[16,17]. Particles were automatically picked *ab initio* using soft-edged disk templates internally generated on *cis*TEM2[16,18]. The picked particle images were boxed, extracted and 2D classified using *cis*TEM2[16]. Selected particles were exported from *cis*TEM2, where global and focused 3D classifications were performed using Relion5[19,20]. The initial 3D reconstructions were carried out *ab initio*, followed by 3D refinement using *cis*TEM2 with refinement of beam tilt and per-particle CTF parameters[16,21]. 3D variability analyses (3DVA) were carried out using cryoSPARC v4[22]. The number of movies and particles used in the final reconstructions are in Table S3.

### Model building

Previous crystal structures of apo (PDB ID: 6EDW) and active state (PDB ID: 6EDZ and 6EE1)[2] were used as initial models and manually rebuilt using COOT[11,23]. All protein models were real-space refined using PHENIX[24], and evaluated using COOT and MolProbity server[25,26]. The cryo-EM maps were deposited in the Electron Microscopy Databank (EMDB), and the coordinates of the atomic models were deposited in Protein Data Bank (PDB)[27,28]. The figures were generated using WARP, Chimera and ChimeraX[29–31].

### Data Deposition

The atomic coordinates and cryo-EM density maps will be available at the PDB and EMDB under accession codes listed in Table S3.

**Results**

***Table S1. Summary of the enzyme kinetics for ICL2 variants in the presence and absence of acetyl-CoA.***

Assays were prepared with 200 nM enzyme, 62.5 µM–16 mM DL-isocitrate, 25 µM acetyl-CoA, 5 mM $MgCl_2$ and 10 mM phenylhydrazine in 50 mM Tris pH 8.0. The reactions were conducted at room temperature (20°C). The concentrations of DL-isocitrate was corrected to the D-enantiomer only, assuming equal ratios of D and L enantiomers, and then used to calculate the kinetic parameters. Data is presented as means ± standard deviations from four independent experiments.

| | **Vmax (µM $min^{-1}$)** | **$K_M$ for isocitrate (mM)** | **$k_{cat}$ ($s^{-1}$)** | **$k_{cat}$/ $K_M$ ($M^{-1}$ $s^{-1}$)** |
|---|---|---|---|---|
| ICL2 | 9.7 ± 0.3 | 1.5 ± 0.2 | 0.8 ± 0.03 | 535 ± 46 |
| ICL2 + acetyl-CoA | 14.6 ± 0.3 | 0.2 ± 0.01 | 1.2 ± 0.02 | 5,106 ± 134 |
| Active variant | 30.0 ± 1.6 | 0.7 ± 0.1 | 2.5 ± 0.1 | 3,480 ± 76 |
| Active variant + acetyl-CoA | 29.1 ± 0.5 | 0.7 ± 0.02 | 2.5 ± 0.04 | 3,389 ± 89 |
| ICL2 Q597E/H598E | 22.5 ± 0.3 | 0.4 ± 0.02 | 1.9 ± 0.03 | 4,742 ± 238 |
| ICL2 Q597E/H598E + acetyl-CoA | 14.4 ± 0.2 | 0.2 ± 0.01 | 1.2 ± 0.02 | 5,398 ± 127 |

***Table S2. Crystallographic statistics for data collection and refinement of the isolated ICL2 C-terminal domain (E640C/N731C/E733C) bound to acetyl-CoA.***
Statistics for the resolution outer shell are shown in parentheses.

| **Data Collection** | **PDB ID 25DG** |
|---|---|
| Space group | $P12_11$ |
| a, b, c (Å) | 37.6, 66.8,131.5 |
| α, β, γ (°) | 90.0, 93.0, 90.0 |
| Beamline | Australian Synchrotron MX2 |
| Wavelength (Å) | 0.95 |
| Resolution Range (Å) | 46.81-2.10 (2.16-2.10) |
| Total Reflections | 530027 (43360) |
| Unique Reflections | 38125 (3084) |
| Multiplicity | 13.9 (14.1) |
| Completeness (%) | 99.8 (99.6) |
| Rpim (within $I$ +/$I$ -) | 0.114 (1.015) |
| Rpim (all $I$ + and $I$ -) | 0.082 (0.730) |
| Mean $I/\sigma(I)$ | 7.4 (1.0) |
| CC1/2 | 0.997 (0.633) |
| **Refinement Statistics** | |
| R factor | 0.232 |
| $R_{free}$ | 0.274 |
| **Structure Validation** | |
| Number of non-H atoms | 5383 |
| *protein* | 4985 |
| *ligands* | 204 |
| *solvent* | 194 |
| Protein residues | 647 |
| RMSZ (bonds) | 0.0034 |
| RMSZ (angles) | 0.774 |
| Ramachandran favoured (%) | 98.7 |
| Ramachandran allowed (%) | 1.3 |
| Ramachandran outliers (%) | 0 |

| | |
|---|---|
| Rotamer outliers (%) | 0 |
| Clash score | 1.37 |
| Wilson B-factor | 30.5 |

***Table S3. Cryo-EM data collection, refinement, and validation statistics.***

| | #1 apo ICL2 | | #2 1 s time-resolved ICL2 | | |
|---|---|---|---|---|---|
| **Data collection and processing** | | | | | |
| Magnification | 165,000 | | 165,000 | | |
| Voltage (kV) | 300 | | 300 | | |
| Electron exposure (e–/Å$^2$) | 60 | | 60 | | |
| Defocus range (µm) | -0.8 to -2.6 | | -0.8 to -2.6 | | |
| Pixel size (Å) | 0.743 | | 0.743 | | |
| Symmetry imposed | C1 | | C1 | | |
| Initial particle images (no.) | 672,154 | | 1,802,410 | | |
| Final particle images (no.) | 279,805 | | 997,105 | | |
| Map resolution (Å) | 3.2 (consensus)<br>3.3 (asymmetric inactive)<br>3.6 (symmetric inactive) | | 3.5 (asymmetric inactive)<br>3.1 (asymmetric active)<br>3.2 (symmetric active)<br>8.0 (active dimerisation) | | |
| FSC threshold | 0.143 | | 0.143 | | |
| Map sharpening *B* factor (Å$^2$) | -90 | | -90 | | |
| **Refinement** | | | | | |
| Initial model used (PDB code) | 6EDW | | 6EDW, 6EDZ, 6EE1 | | |
| PDB code | 12HU | 12HV | 12IB | 12IA | 12IB |
| Model composition | #1 asymmetric inactive | #1 symmetric inactive | #2 asymmetric inactive | #2 asymmetric active | #2 symmetric active |

| | | | | | |
|---|---|---|---|---|---|
| Non-hydrogen atoms | 17353 | 17376 | 17044 | 20104 | 22893 |
| Protein residues | 2222 | 2226 | 2181 | 2550 | 2878 |
| Nucleic acid residues | 0 | 0 | 0 | 0 | 0 |
| Ligands | 0 | 0 | 0 | 2 | 4 |
| Refinement (Phenix) | | | | | |
| Map correlation coefficient (whole unit cell) | 0.59 | 0.50 | 0.60 | 0.69 | 0.61 |
| Map correlation coefficient (around atoms) | 0.80 | 0.72 | 0.82 | 0.89 | 0.86 |
| R.m.s. deviations | | | | | |
| Bond lengths (Å) | 0.005 | 0.006 | 0.007 | 0.007 | 0.006 |
| Bond angles (°) | 0.739 | 0.792 | 0.938 | 0.853 | 0.824 |
| Validation | | | | | |
| MolProbity score | 1.81 | 1.85 | 1.90 | 1.88 | 1.76 |
| Clashscore | 13.3 | 16.0 | 18.4 | 17.7 | 13.5 |
| Poor rotamers (%) | 0.11 | 0.23 | 0.00 | 0.05 | 0.00 |
| Ramachandran plot | | | | | |
| Favored (%) | 96.98 | 97.22 | 97.29 | 97.33 | 97.35 |
| Allowed (%) | 2.98 | 2.73 | 2.71 | 2.67 | 2.65 |
| Disallowed (%) | 0.05 | 0.05 | 0.00 | 0.00 | 0.00 |

| | #3 0.15 s time-resolved ICL2 | #4 30 min ICL2 |
|---|---|---|
| **Data collection and processing** | | |
| Magnification | 81,000 | 105,000 |
| Voltage (kV) | 300 | 300 |
| Electron exposure (e–/Å$^2$) | 60 | 60 |

| | | | |
|---|---|---|---|
| Defocus range (μm) | -0.8 to -2.6 | -0.8 to -2.6 | |
| Pixel size (Å) | 0.857 | 0.827 | |
| Symmetry imposed | C1 | C1 | |
| Initial particle images (no.) | 821,924 | 1,351,602 | |
| Final particle images (no.) | 146,216 | 932,554 | |
| Map resolution (Å) | 4.0 | 2.9 (asymmetric active) | |
| | | 2.9 (symmetric active) | |
| FSC threshold | 0.143 | 0.143 | |
| Map sharpening *B* factor ($Å^2$) | -90 | -90 | |
| **Refinement** | | | |
| Initial model used (PDB code) | 6EDW | 6EDZ, 6EE1 | |
| PDB code | 12HY | 12HX | 12HW |
| Model composition | | #4 asymmetric active | #4 symmetric active |
| Non-hydrogen atoms | 18045 | 20137 | 22700 |
| Protein residues | 2313 | 2554 | 2855 |
| Nucleic acid residues | 0 | 0 | 0 |
| Ligands | 0 | 2 | 4 |
| Refinement (Phenix) | | | |
| Map correlation coefficient (whole unit cell) | 0.65 | 0.65 | 0.63 |
| Map correlation coefficient (around atoms) | 0.66 | 0.88 | 0.88 |
| R.m.s. deviations | | | |
| Bond lengths (Å) | 0.007 | 0.004 | 0.006 |
| Bond angles (°) | 1.15 | 0.721 | 0.905 |
| Validation | | | |

| | | | |
|---|---|---|---|
| MolProbity score | 1.6 | 1.47 | 1.74 |
| Clashscore | 12.22 | 8.76 | 17.27 |
| Poor rotamers (%) | 0.22 | 0 | 0.44 |
| Ramachandran plot | | | |
| Favored (%) | 98.7 | 98.65 | 98.35 |
| Allowed (%) | 1.3 | 1.35 | 1.61 |
| Disallowed (%) | 0 | 0 | 0.04 |

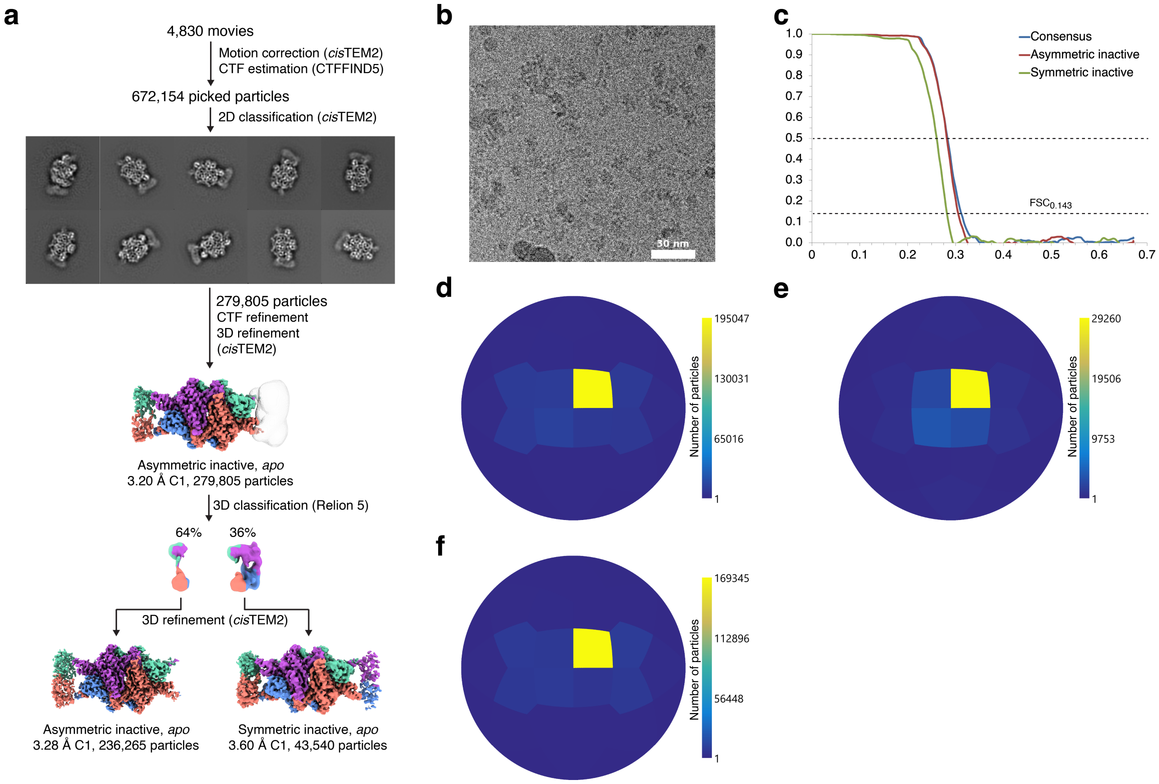


***Figure S1. Cryo-EM data processing workflow of apo ICL2.***

**a,** Cryo-EM data processing workflow of apo ICL2. In the apo state, ICL2 exists in partially and fully inactive states. **b,** A representative micrograph of apo ICL2. The scale bar is 50 nm. **c,** Half-map Fourier shell correlation (FSC) plots. Dotted lines indicate $FSC_{0.5}$ and $FSC_{0.413}$. **d,** Angular distribution plot of the consensus reconstruction. **e,** Angular distribution plot of the fully inactive ICL2. **f,** Angular distribution plot of the partially inactive ICL2.

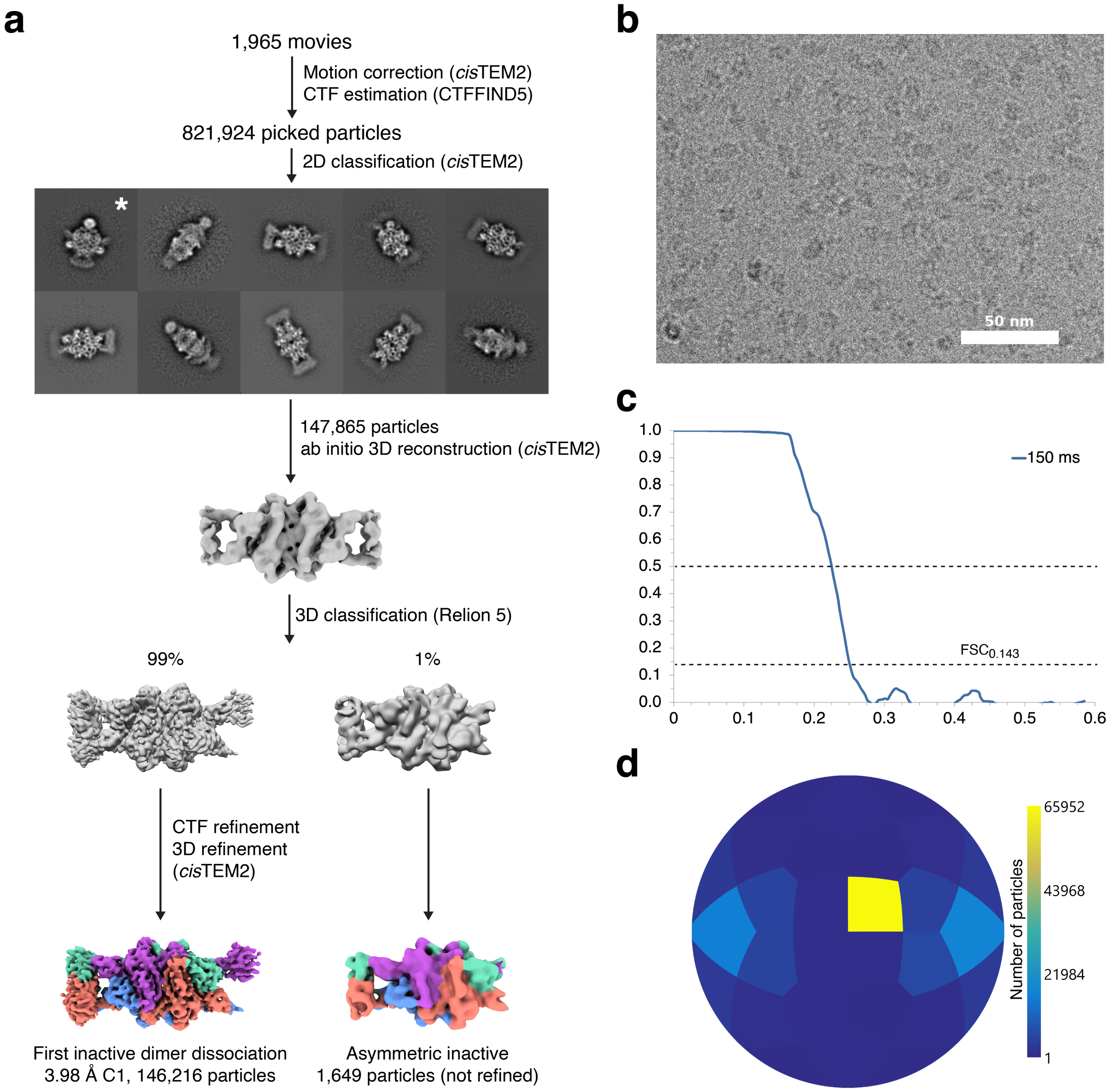


***Figure S2. 0.15 s time-resolved cryo-EM data processing workflow.***
**a,** 0.15 s time-resolved cryo-EM data processing workflow of ICL2. With 0.15 s sample mixing time, ICL2 predominantly adopts an inactive state where one C-terminal dimer has dissociated, with density observed for only one monomeric C-terminal domain. **b,** A representative micrograph of ICL2 with 0.15 s mixing time. The scale bar is 50 nm. **c,** Half-map Fourier shell correlation (FSC) plot of the first inactive C-terminal dimer dissociation. Dotted lines indicate $FSC_{0.5}$ and $FSC_{0.413}$. **d,** Angular distribution plot of the first inactive C-terminal dimer dissociation.

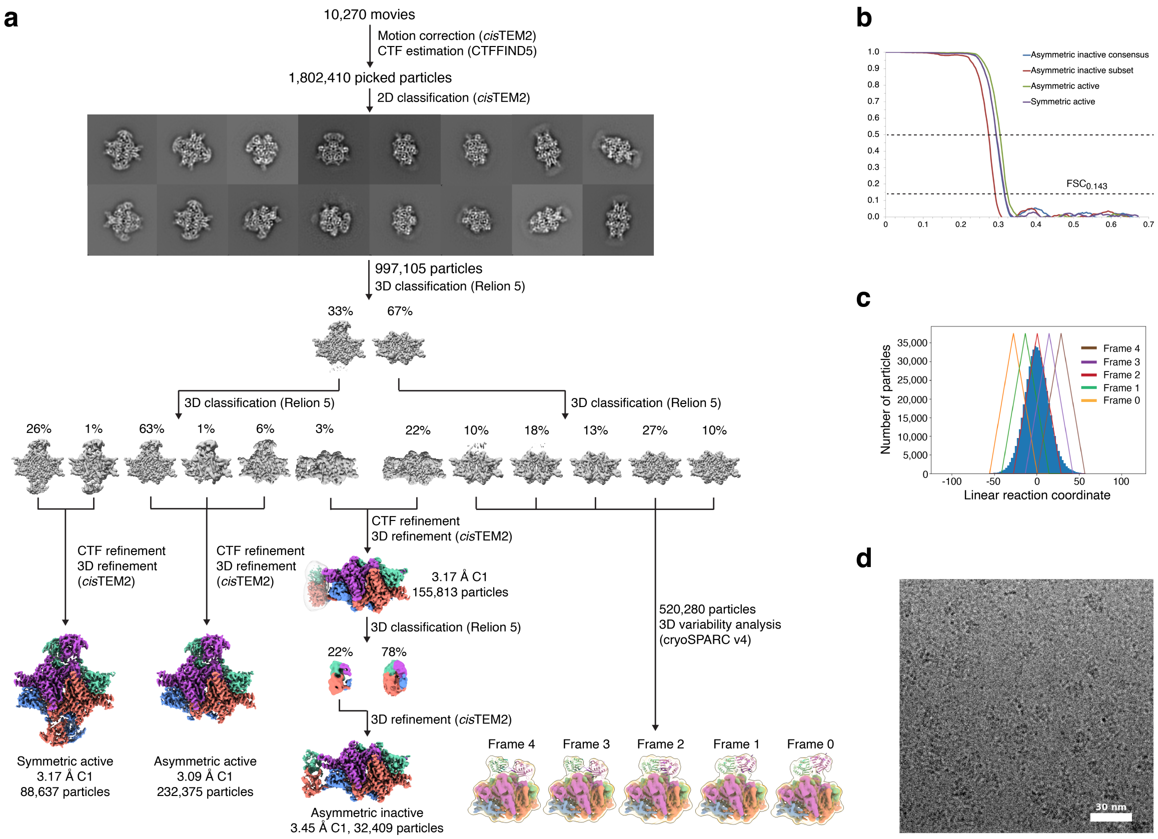


***Figure S3. 1 s time-resolved cryo-EM data processing workflow***

**a,** 1 s time-resolved cryo-EM data processing workflow of ICL2. With 1 s sample mixing time, ICL2 exists in a spectrum of different states along the activation pathway. **b,** Half-map Fourier shell correlation (FSC) plots. Dotted lines indicate $FSC_{0.5}$ and $FSC_{0.413}$. **c,** 3D variability analysis showing fractionation of the particles over the linear reaction coordinate in five overlapping windows. **d,** A representative micrograph of ICL2 with 0.15 s mixing time. The scale bar in 50 nm.

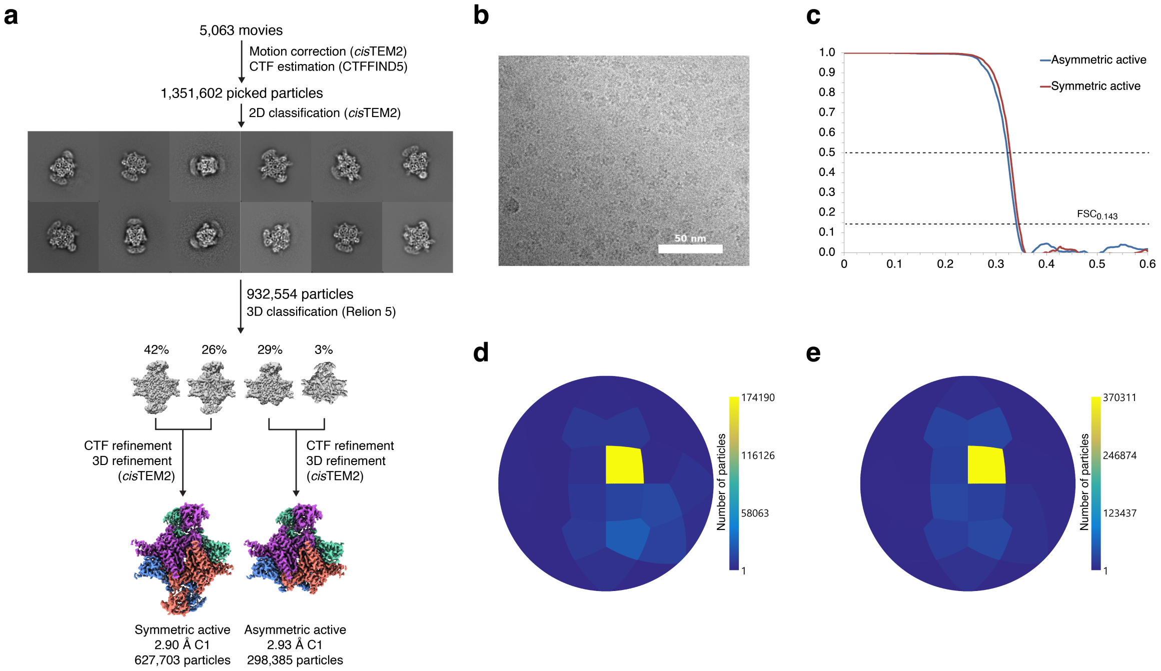


***Figure S4. 30 min time-resolved cryo-EM data processing workflow***

**a,** 30 min time-resolved cryo-EM data processing workflow of ICL2. With 30 min mixing time, ICL2 exists in partially or fully active states. **b,** A representative micrograph of ICL2 with 30 min mixing time. The scale bar is 50 nm. **c,** Half-map Fourier shell correlation (FSC) plots. Dotted lines indicate $FSC_{0.5}$ and $FSC_{0.413}$. **d,** Angular distribution plot of the partially active state reconstruction. **e,** Angular distribution plot of the fully active state reconstruction.

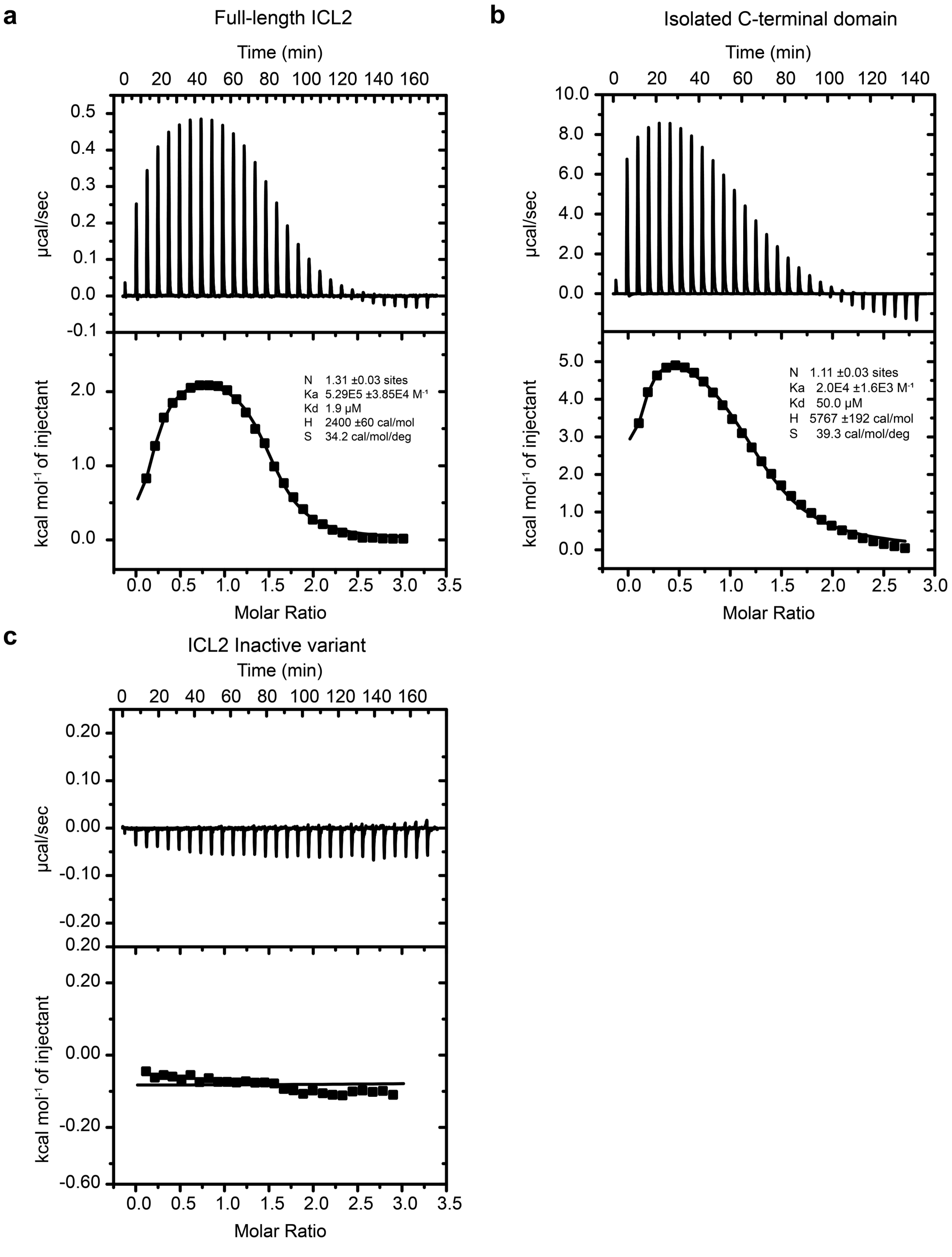


***Figure S5. Comparison of acetyl-CoA binding to the full-length ICL2, the isolated C-terminal domain, and to the inactive variant determined by isothermal titration calorimetry.***

**a,** 1000 μM Acetyl-CoA was titrated against 60 μM full-length ICL2 **b,** 7000 μM acetyl-CoA was titrated against 550 μM of the isolated C-terminal domain (residues 601-766) **c,** 500 μM Acetyl-CoA was titrated against 50 μM full-length inactive variant

(D377C/L601C, F381C/T604C) with engineered disulfide bonds between the linker and the helical subdomain in the inactive conformation. The upper panel shows the raw titration data, and the lower panel shows the integrated heats of binding. The solid line represents the best fit curve. Assays were undertaken in a buffer containing 10 mM phosphate buffer pH 8.0 and 150 mM NaCl. Errors represent the fitting errors.

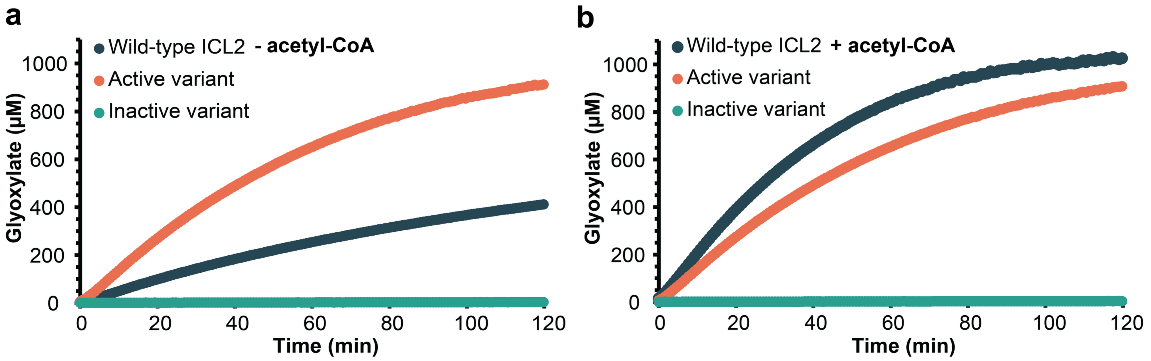


***Figure S6. Comparison of reaction time course of ICL2 and engineered active and inactive variants with and without acetyl-CoA.***

Assays were prepared with 200 nM enzyme, 1 mM DL-isocitrate, 5 mM $MgCl_2$, 150 mM NaCl, and 10 mM phenylhydrazine in 50 mM Tris pH 8.0. Reactions were conducted at room temperature (20°C). **a,** Assays were undertaken in the absence of acetyl-CoA with wild-type ICL2, the active variant (E640C/N731C/E733C), and the inactive variant (D377C/ L601C,F381C/T604C). **b,** 25 µM acetyl-CoA was pre-incubated with the enzymes before preparing the assay. Data are presented as means ± standard deviations from four independent experiments.

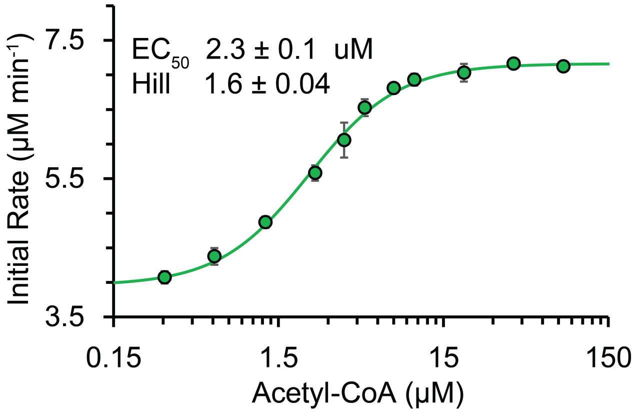


***Figure S7. Acetyl-CoA binding to Mtb-ICL2 shows positive cooperativity.***

Assays were prepared with 200 nM ICL2, 3 mM DL-isocitrate, 0-80 µM acetyl-CoA, 5 mM $MgCl_2$, 150 mM NaCl, and 10 mM phenylhydrazine in 50 mM Tris pH 8.0. Reaction temperature was room temperature (20 °C). Data are presented as means ± standard deviations from four independent experiments.

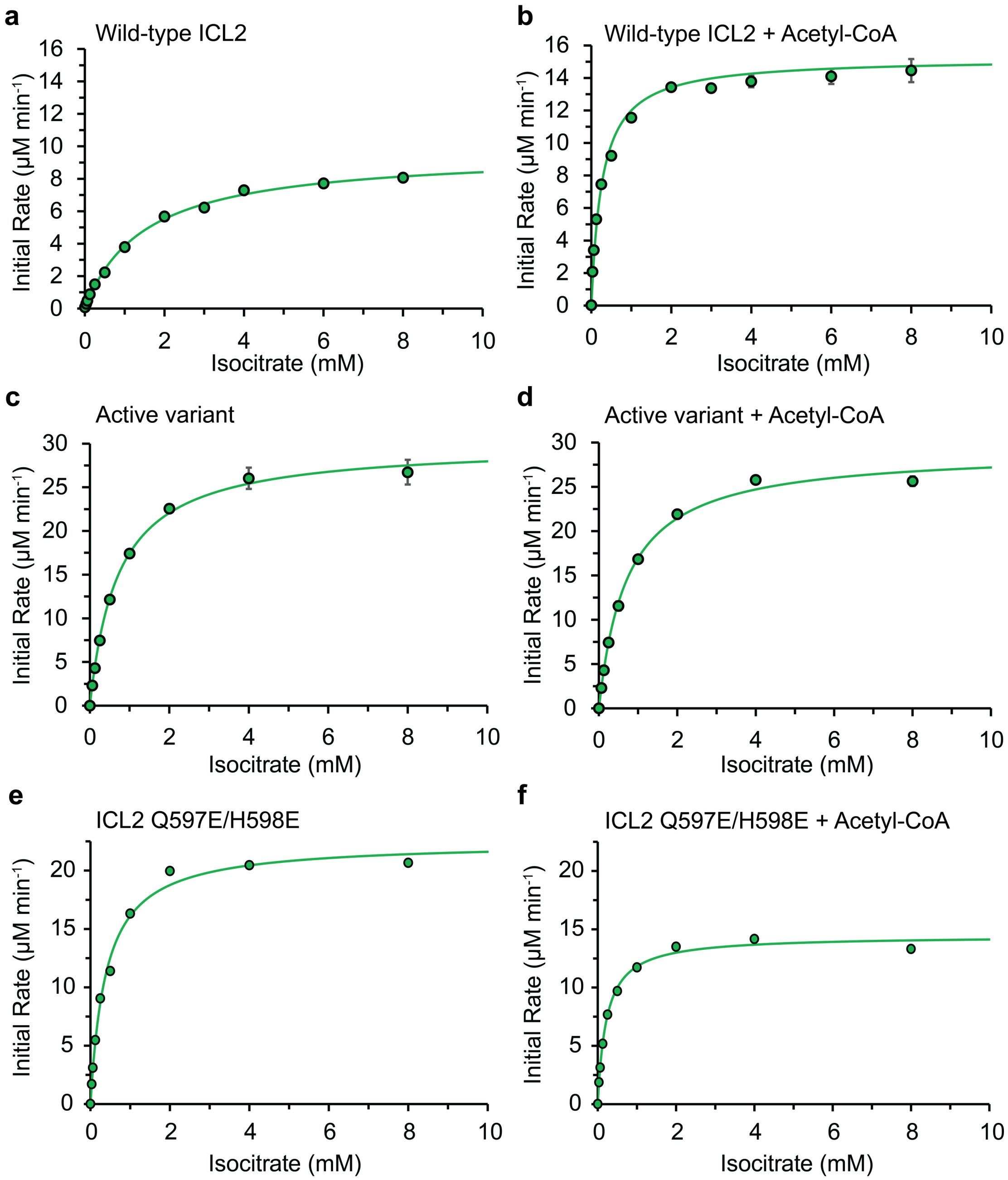


***Figure S8. Michaelis-Menten kinetics of ICL2 variants for DL-isocitrate in the absence and presence of acetyl-CoA.***

**a-b,** The $K_M$ value for wild-type ICL2 in the presence and absence of 25 µM acetyl-CoA. **c-d,** The $K_M$ value for the active variant (E640C/N731C/E733C) in the presence and absence of 25 µM acetyl-CoA. **e-f**. The $K_M$ value for ICL2 Q597E/H598E in the presence and absence of 25 µM acetyl-CoA. Assays were prepared with 200 nM enzyme, 62.5 µM–16 mM DL-isocitrate, 5 mM $MgCl_2$, 150 mM NaCl, and 10 mM phenylhydrazine in 50 mM Tris pH 8.0. Reactions were conducted at room temperature (20°C). The concentration of DL-isocitrate was corrected to the D-enantiomer only, assuming equal ratios of D and L enantiomers, and then used to calculate the kinetic parameters. Data are presented as means ± standard deviations from four independent experiments.

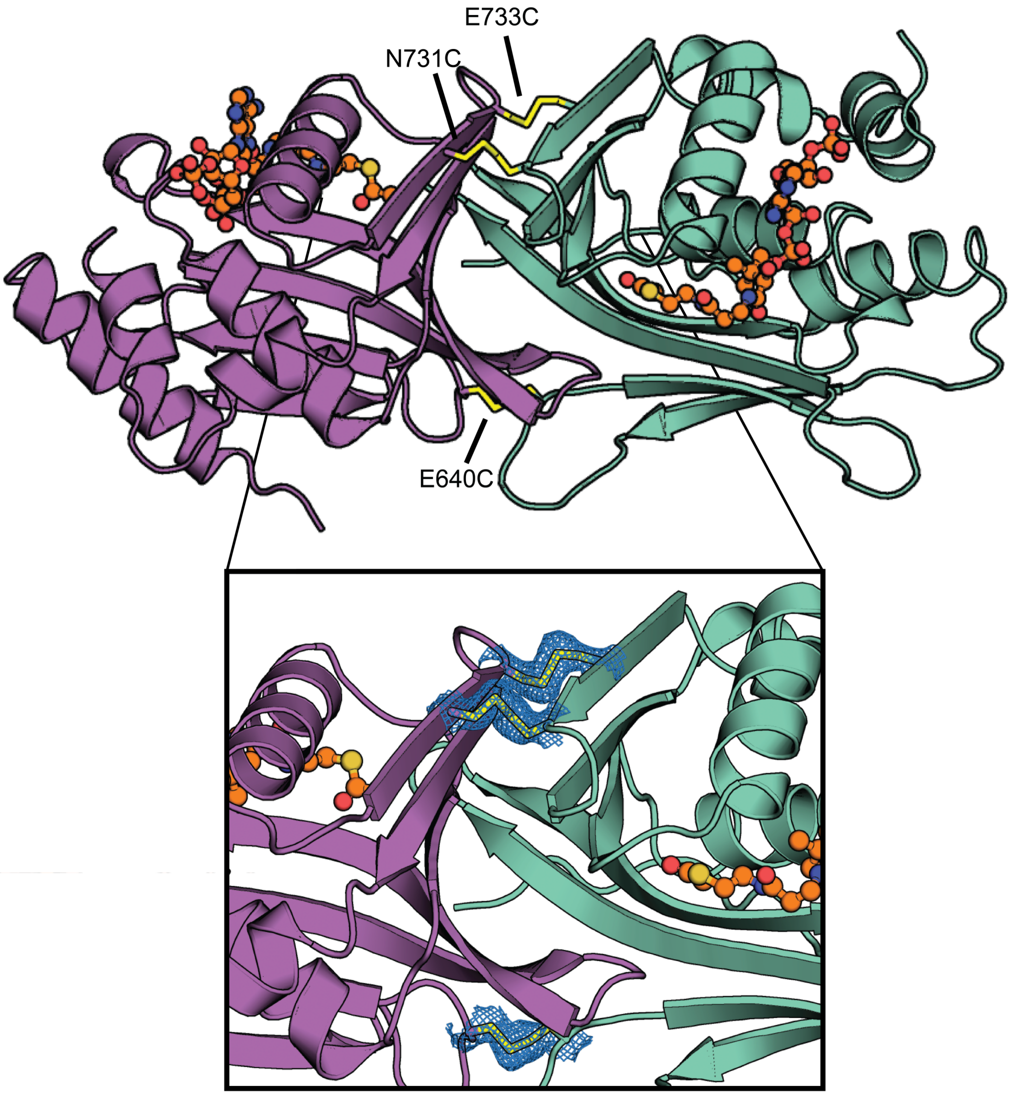


***Figure S9. The co-crystal structure of the isolated, active C-terminal domain bound to acetyl-CoA.***

The crystal structure of isolated C-terminal domain (E640C/N731C/E733C) showing the C-terminal domains form a dimer with disulfide bonds formed between the mutated residues E640C, N731C, and E733C from both monomers. Acetyl-CoA (orange ball-and-stick model) is bound in the ligand-binding pocket. The inset shows the $2F_o$–$F_c$ electron density for the disulfide bonds with $\sigma = 1$.

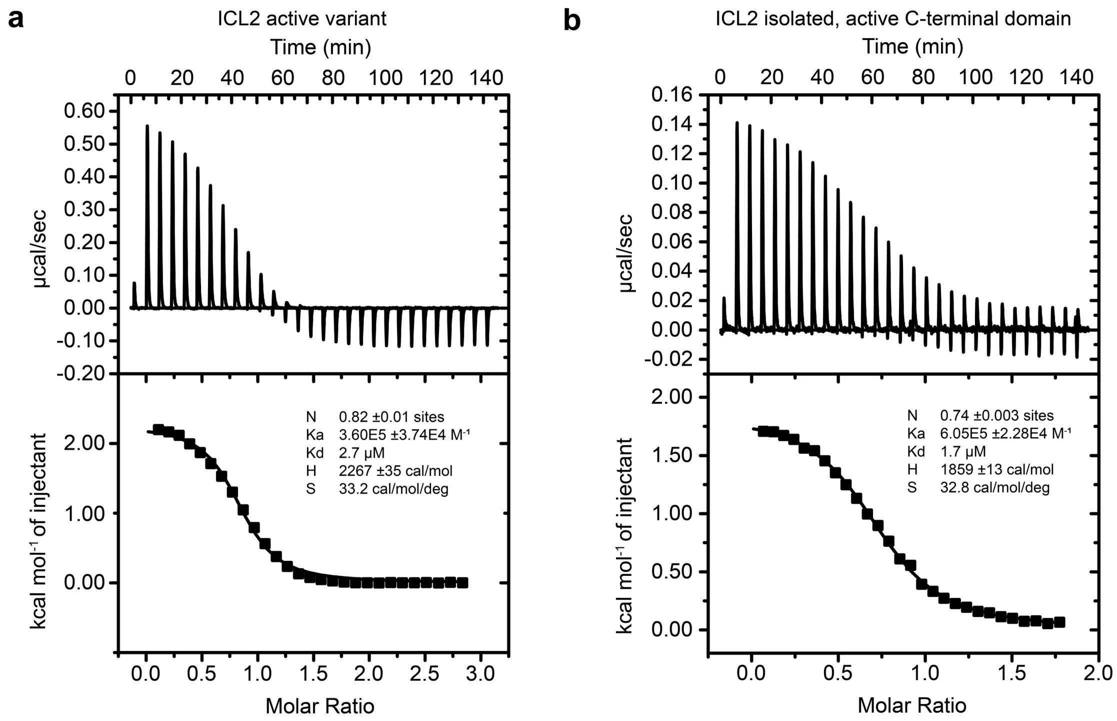


***Figure S10. ITC analysis of acetyl-CoA binding to the full-length active variant and the isolated, active C-terminal domain.***

**a**, 500 µM Acetyl-CoA was titrated against 50 µM of the full-length active ICL2 (E640C/N731C/E733C). b, 250 µM Acetyl-CoA was titrated against 30 µM of the isolated, active C-terminal domain. The upper panel shows the raw titration data, and the lower panel shows the integrated heats of binding. The solid line represents the best fit curve. Assays were undertaken in a buffer containing 10 mM phosphate buffer pH 8.0 and 150 mM NaCl. Errors represent the fitting errors.

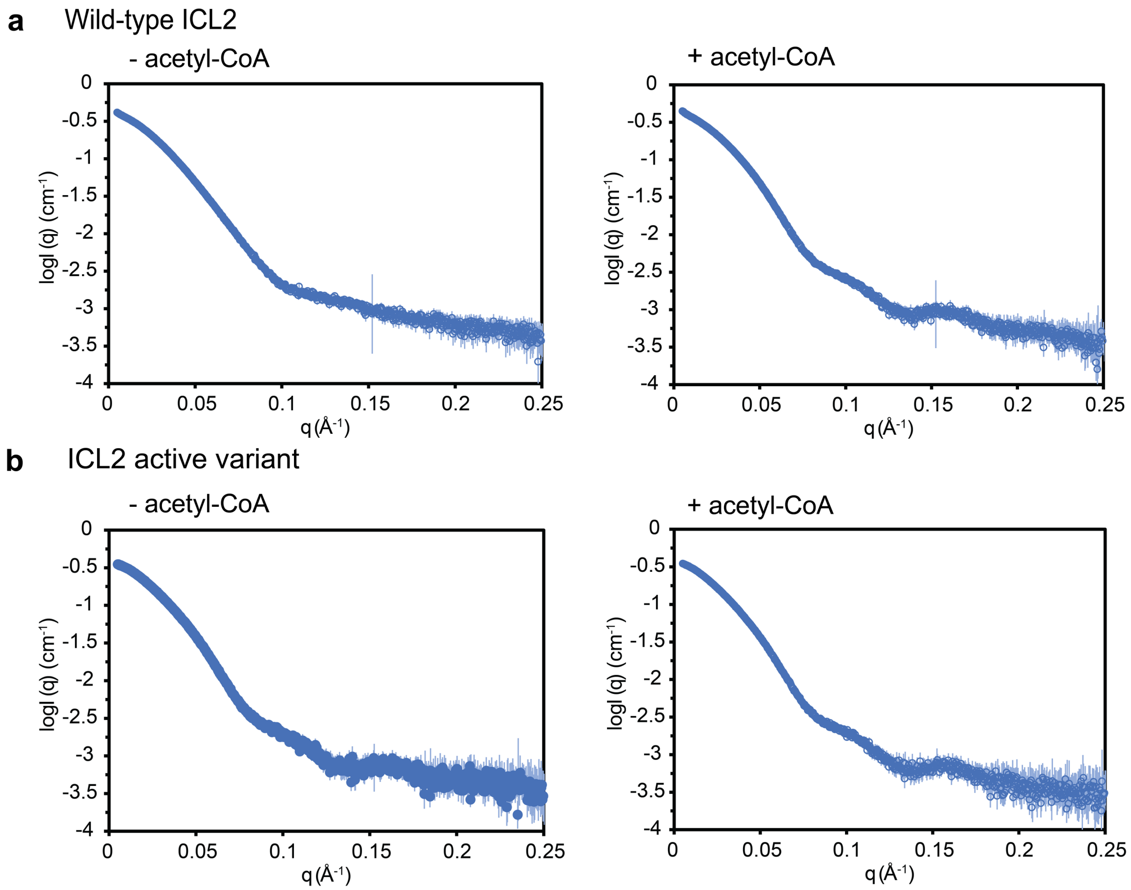


**Figure S11. Small-angle X-ray scattering of wild-type ICL2 and the active2 variants in the presence and absence of acetyl-CoA.**

**a,** Scattering of wild-type ICL2 in the presence and absence of acetyl-CoA, indicating a considerable difference in ICL2 conformation in these two states. **b,** The active variant (E640C/N731C/E733C) shows a similar scattering profile to the wild-type ICL2, even in the absence of acetyl-CoA. ICL2 constructs were extensively dialysed against 20 mM Tris pH 8.0, 150 mM NaCl, 5% glycerol (v/v). Proteins were diluted to 2 mg/mL and, where appropriate, incubated with 500 µM acetyl-CoA before SAXS analysis.

### Acknowledgements

We thank Drs. Daniel Smith and Andrew Woehler at Janelia Experimental Technology (jET) and the members of HHMI Janelia Research Campus Cryo-EM Shared Resources Facility for helpful discussion and advice. We thank Dr. Stephen P. Muench and Isobel J. Hirst (University of Leeds) for expert advice and support on the time-resolved microfluidic mixer-sprayer instrument.
